# Individual Differences Elucidate the Perceptual Benefits Associated with Robust Temporal Fine-Structure Processing

**DOI:** 10.1101/2023.09.20.558670

**Authors:** Agudemu Borjigin, Hari M. Bharadwaj

## Abstract

The auditory system is unique among sensory systems in its ability to phase lock to and precisely follow very fast cycle-by-cycle fluctuations in the phase of sound-driven cochlear vibrations. Yet, the perceptual role of this temporal fine structure (TFS) code is debated. This fundamental gap is attributable to our inability to experimentally manipulate TFS cues without altering other perceptually relevant cues. Here, we circumnavigated this limitation by leveraging individual differences across 200 participants to systematically compare variations in TFS sensitivity to performance in a range of speech perception tasks. TFS sensitivity was assessed through detection of interaural time/phase differences, while speech perception was evaluated by word identification under noise interference. Results suggest that greater TFS sensitivity is not associated with greater masking release from fundamental-frequency or spatial cues, but appears to contribute to resilience against the effects of reverberation. We also found that greater TFS sensitivity is associated with faster response times, indicating reduced listening effort. These findings highlight the perceptual significance of TFS coding for everyday hearing.

**Significance Statement:** Neural phase-locking to fast temporal fluctuations in sounds–temporal fine structure (TFS) in particular– is a unique mechanism by which acoustic information is encoded by the auditory system. However, despite decades of intensive research, the perceptual relevance of this metabolically expensive mechanism, especially in challenging listening settings, is debated. Here, we leveraged an individual-difference approach to circumnavigate the limitations plaguing conventional approaches and found that robust TFS sensitivity is associated with greater resilience against the effects of reverberation and is associated with reduced listening effort for speech understanding in noise.

---

Human connection and communication fundamentally rely on the auditory system’s capacity to encode and process complex sounds such as speech and music. Regardless of complexity, all acoustic information we receive from our environment is conveyed through the firing rate and spike timing of cochlear neurons (i.e., rate-place vs. temporal coding) (1). Temporal information in sound-driven cochlear responses is comprised of two components: rapid variations in phase—the temporal fine structure (TFS), and slower amplitude variations —the temporal envelope (2). Neurons in the auditory system can robustly track both TFS (3) and envelope (4) through phase-locked firing. Strikingly, neural phase locking to TFS extends at least up to 1400 Hz in the peripheral auditory system (5–7), a feat unmatched by other sensory modalities. In comparison, phase-locked information in the visual and somatosensory systems extends only to about 50 Hz (8, 9). However, this uniquely high upper-frequency limit of phase locking in the auditory system only exists at the peripheral level (5, 10). Along the ascending pathway, the phased-locked temporal code appears to be progressively transformed into a rate-place representation (11). It seems that the auditory system initially invests heavily in this exquisite and metabolically expensive (12, 13) phaselocked temporal code but then “repackages” the code into a different form for downstream processing. How this initial neural coding of TFS ultimately contributes to perception, and if and how its degradation leads to perceptual deficits is a fundamental open question not only for the neuroscience of audition, but also for clinical audiology. Yet, the significance of this peripheral TFS phase-locking in the auditory system remains controversial (5, 14–20).

Psychophysical experiments in *quiet* sound booths suggest that TFS may play a role in sound localization (21, 22) and pitch perception (through fundamental-frequency or F0 cues) (23–25). Both spatial and F0 information can serve as primary cues for target-background segregation and selective attention in more realistic listening settings, yielding a masking release of about 5 dB each (26–34). Yet, whether this masking release is attributable to TFS coding is debated. This is because the other component of sound—the temporal envelope, despite eliciting weaker pitch or spatial percepts in quiet, can provide a similar degree of masking-release benefit in noise (17, 35). Furthermore, TFS-based spatial cues are more susceptible to corruption from reverberation than envelope-based spatial cues (36, 37) by virtue of being perceptually dominant primarily at low-frequencies up to about 1400 Hz (7, 38), where reverberation is more pronounced (37). Thus, despite many decades of intensive research, whether phase-locked temporal coding of TFS would introduce additional masking-release benefits in reverberant listening conditions remains unclear.

A key challenge to understanding the perceptual role of TFS phase locking is that sub-band vocoding, which is the most common technique employed to investigate this question, is inherently limited (21, 39–43). Vocoding has been used to acoustically dissociate TFS from envelope by creating stimuli with a constant envelope (i.e., sub-band amplitude) while manipulating the TFS (i.e., sub-band phase). Unfortunately, this clean dissociation at the acoustic level is not maintained at the output of cochlear processing, which inter-converts some of the TFS cues to amplitude fluctuations (16, 44, 45). Recent approaches to investigate the perceptual significance of TFS coding have leveraged deep neural networks (DNN) and evaluated how the performance of DNNs trained on a range of tasks is affected when TFS cues are degraded in the input (19, 46). However, similarly to perceptual studies that employ sub-band vocoding, the DNN studies are also subject to the introduction of confounding effects at the output of cochlear processing. Some studies, such as Hopkins et al., 2008 (39) and Smith et al., 2002 (21) have employed stimuli that combine envelope and TFS information from distinct speech utterances to study the role of TFS. However, these studies are subject to a broader limitation of stimulus-manipulation approaches: participants may use and weight TFS cues differently depending on the availability of other redundant cues, and thus differently in synthetic versus naturalistic stimuli.

An alternative approach that can overcome these limitations is to avoid any stimulus manipulations, but directly measure individual differences in TFS processing and compare them to individual differences in speech-in-noise outcomes tested with intact, minimally-processed stimuli. The individual differences approach has been successfully used to address other fundamental questions in the neuroscience of audition (47–50). At the time of of this study, the individual-difference approach has not been used to explore the role of TFS for speech-in-noise perception, as robust individual-level measures were only recently established by comparing both EEG and behavioral measures of TFS coding (51). Since, however, Vinay and Moore, 2023 (52) have used a similar approach to examine the role of TFS and place coding for frequency discrimination task at 2 kHz.

Here, we leveraged individualized TFS processing measures developed in our previous work, and adapted them for remote testing to circumnavigate the COVID-19-related restrictions (53). We hypothesized that TFS plays an important role in everyday hearing. To elucidate the role of TFS in everyday listening, we compared individual TFS sensitivity to individual participants’ speech-perception outcomes under various types of noise interference. The speech-in-noise test battery included ten different listening conditions, representing many important aspects of everyday listening where TFS phase locking has conventionally been thought to play a role. We predicted that individuals with better TFS sensitivity would benefit more from F0 and spatial cues in noisy listening settings because of the hypothesized role of TFS in pitch perception and sound localization (21–25). Because reverberation impairs TFS-based spatial cues (36) and spatial selective attention (54), we predicted that individuals with better TFS sensitivity would be less affected by reverberation.

Lastly, we hypothesized that individuals with better TFS sensitivity would expend less listening effort and show more release from informational masking. Informational masking occurs when listeners fail to segregate or select the target sound components in the mixture despite minimal direct spectrotemporal overlap between the target and maskers. Both listening effort and listening under conditions of informational masking have been linked to a number of central auditory and cognitive processes (55–57); the availability of robust TFS cues is thought to be beneficial to these processes (58–60). There is now considerable literature suggesting that performance scores alone do not capture the widely varying degree of cognitive effort that different participants have to put in to reach the same score. Response times have thus found increasing use in the “listening effort” literature as a measure that is sensitive to differences in the cognitive burden experienced by different participants (61, 62). Accordingly, we measured response times in addition to speech-in-noise scores. The automated and parallel nature of the online measurements allowed us to rapidly collect data from a large cohort of 200 participants, affirming the promise and advantages of online behavioral psychoacoustical studies (50, 53). Figure 1 illustrates the design of this study. The results revealed that better TFS processing, although not associated with greater masking release [confirming the results from Füllgrabe et al., 2015 (63)], provided resilience against reverberation, and lessened listening effort. Given that reverberation is a common source of signal corruption in everyday listening, and that listening effort is often a primary patient complaint in the audiology clinic, these findings highlight the perceptual significance of TFS coding in everyday communication.

**Fig. 1.**
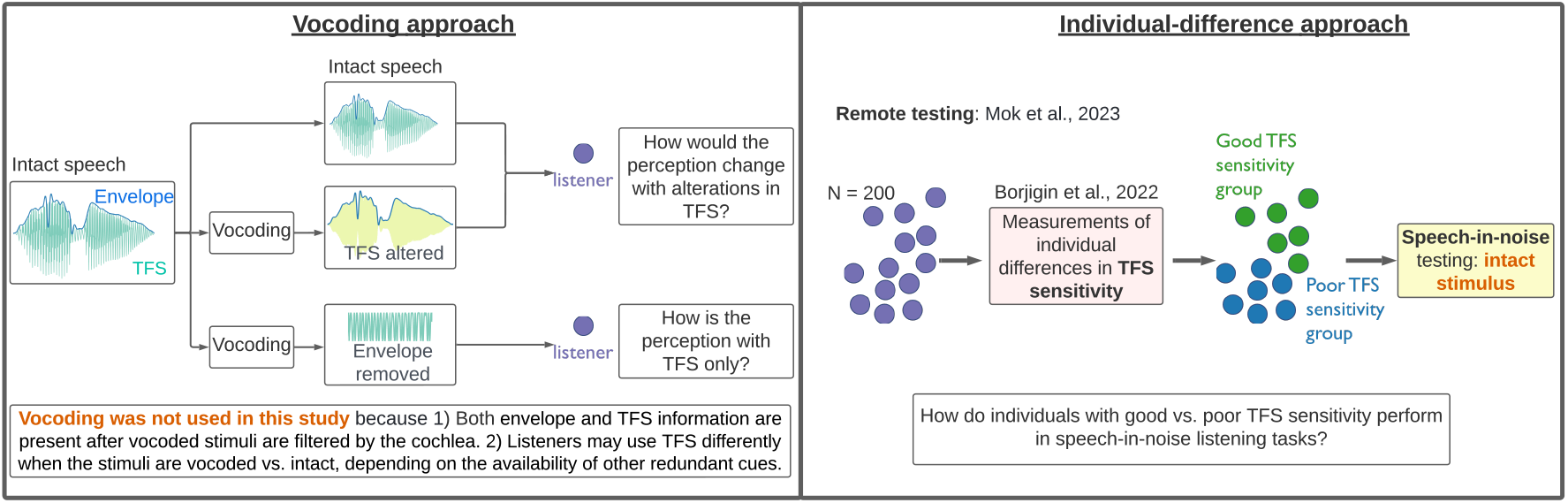
Contrasting the conventional vocoding approach (left) for studying TFS with the individual-difference approach adopted in this study (right).

## Results

### Binaural temporal sensitivity measures captured individual differences in TFS processing fidelity

Figure 2 (a) is a scatter plot of the individual differences that we observed for our two binaural temporal sensitivity measures—ITD discrimination and binaural FM detection (FM of opposite phase in the two ears). Metrics of individual TFS sensitivity commonly used in the literature are prone to the impact of extraneous “non-sensory” variables (51) such as attention and motivation. Here, ILD discrimination was used as a surrogate measure to control for “non-sensory” factors as well as as aspects of binaural hearing unrelated to the basic TFS code. These TFS metrics were accordingly “adjusted” by regressing out the ILD sensitivity scores from each measure. The individual differences in these “adjusted” TFS metrics are more likely driven by true individual differences in TFS processing (See Methods for further details). Individual ILD sensitivity data are shown in Figure 2 (b), which also indicates substantial individual variability. Note that in Figure 2 (a), the TFS metrics are shown after regressing out the ILD measure, and vice versa.

**Fig. 2.**
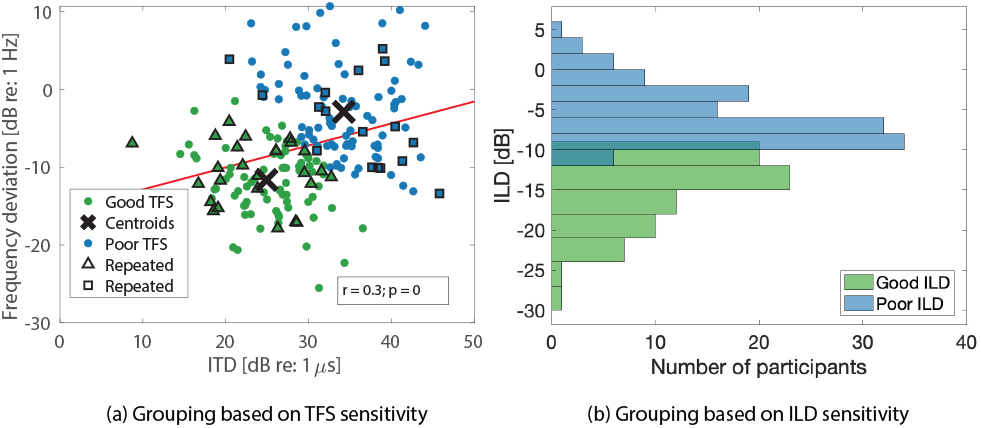
(a) An illustration of cluster assignment based on the combination of ITD and binaural FM thresholds. Note that the marked participants (triangles and squares) returned for replication measurements (see Figure 7 showing replication data). (b) Group assignment based on ILD sensitivity. Note that the ITD, ILD, binaural FM sensitivity values displayed are the residuals after regression.

The adjusted binaural FM detection and ITD discrimination measures were significantly correlated (*r* = 0.3, *p <*.0001) indicating a common underlying source of variance attributable to TFS processing. Accordingly, participants were divided into two groups by a clustering algorithm based on these two measures into “Good-” vs “Poor-TFS” groups. As can be expected from the grouping procedure, Figures 3 (a) and (b) show a clear separation of the two groups’ psychometric curves in the ITD and binaural FM measurements, respectively. More importantly, when the ILD data, which were not used for grouping, were plotted for these two groups, there was no separation in the psychometric curves [Figure 3 (c)], demonstrating that the groupings are orthogonal. The construction of groups based on common variance across the TFS measures after eliminating common variance with ILD sensitivity ensures that the grouping in Figure 2 (a) is mainly based on individuals’ TFS sensitivity, rather than other unrelated factors. Note also that there is no significant difference in age between two groups (“Good TFS” group: mean age of 30.4 years with an std of 7.7 years; “Poor TFS” group: mean age of 32.1 years with an std of 8.4 years). To corroborate this further, individuals were also grouped based on their ILD discrimination thresholds, as shown in Figure 2 (b). For this alternative grouping, a clear separation is evident in the psychometric curves for ILD discrimination [Figure 3 (f)] (as expected), but not in the curves for TFS measurements [Figure 3 (d) or (e)], consistent with the notion that the grouping in Figure 2 (b) captures “non-TFS” variability instead of TFS sensitivity. This alternative non-TFS regrouping of participants is used as a control in the experiments probing the association between TFS processing and speech-in-noise outcomes.

**Fig. 3.**
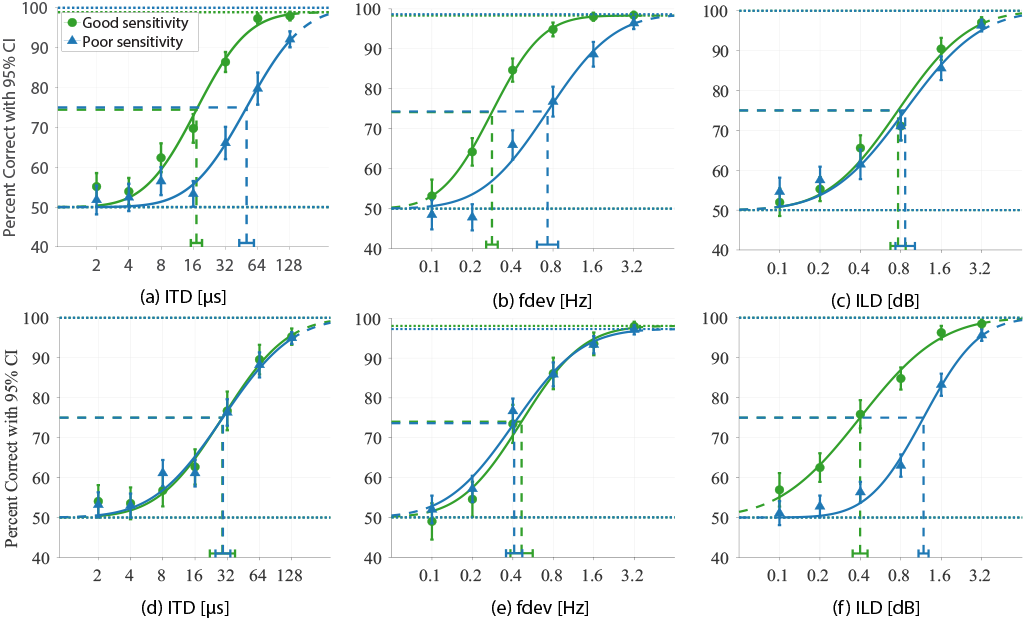
(a)-(c): Psychometric curves for Good vs. Poor TFS-sensitivity groups. Left: ITD; mid: binaural FM (fdev: frequency deviation); right: ILD. (d)-(f): Psychometric curves for Good vs. Poor ILD-sensitivity groups. Error bars represent within-group standard error of the mean.

The web-based measurements in the present study produced data that were comparable to the data not only from our previous in-person study, but also from other labs. Figure 4 shows comparisons for FM detection and ITD discrimination measurements across studies. The left panel compares online measurement of binaural FM detection with in-person results from (64–67). The right panel includes a sample of in-person studies that measured ITD discrimination (7, 51, 68, 69). (51) is our previous in-person study. These results further validate our choice of TFS sensitivity measures.

**Fig. 4.**
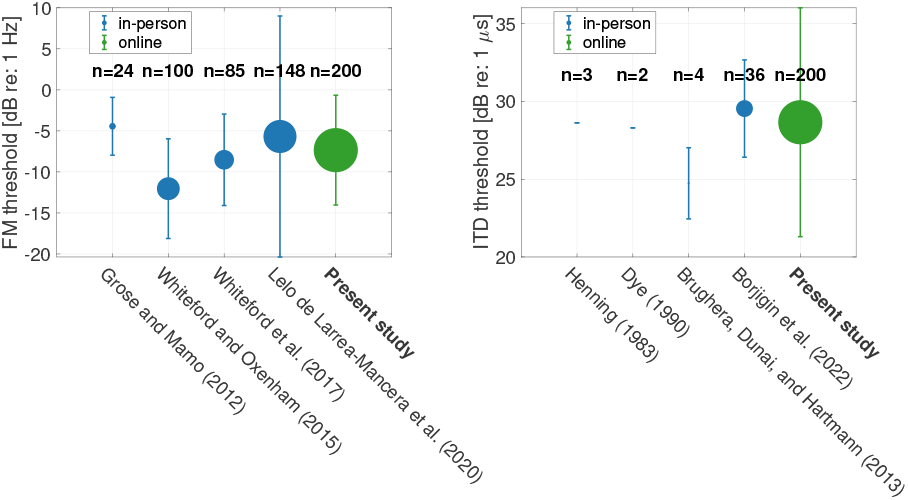
A sample of published reports of binaural FM detection thresholds (left) and ITD discrimination threshold (right) for comparison with the present study. Error bars represent +/-1 std. The std of ITD detection thresholds could not be determined for (68) and (69). The size of the dot represents the number of participants.

### Better TFS sensitivity is not associated with additional masking-release benefit

To understand the functional role of TFS in everyday hearing, we measured participants’ speech intelligibility under various types of noise interference, in addition to evaluating TFS sensitivity. Rather than absolute speech reception threshold (SRT, the lowest/noisiest level at which a person can understand speech in noise), Figure 5 (a) depicts the masking release. Masking release refers to improved noise tolerance associated with the following cues: F0 difference between the target and background speakers, spatial separation between the target and maskers, combination of F0 and spatial cues, and finally when the background noise was non-speech stationary noise instead of speech babble. The masking release effects observed in this study are consistent with those reported in previous research: 1) With F0 separation, the participants could more easily identify the target compared to when the target and background had similar F0 (i.e., the reference condition). This F0-based masking release was about 5 dB (Good TFS group: mean = 4.8, std = 0.5; Poor TFS group: mean = 5.3, std = 0.4), which matches previous reports from (32–34). 2) The masking release was around 3 dB when the target and background were spatially separated (Good TFS group: mean = 2.9, std = 0.4; Poor TFS group: mean= 3.7, std = 2.3), which again aligns with earlier reports (26–31). 3) When both F0 and spatial cues were available, the masking-release benefits appeared to be cumulative, totaling about 10 dB as demonstrated in the “F0 + space” condition (Good TFS group: mean = 10.3, std = 0.8; Poor TFS group: mean = 9.5, std = 0.6). Indeed, it has previously been shown that F0 differences aid participants in spatially separating competing sounds (70). 4) A masking release of about 19 dB was observed when the background noise was switched from 4-talker babble to non-speech stationary noise, as shown in the “steady noise” condition (Good TFS group: mean = 19.2, std = 0.7; Poor TFS group: mean = 18, std = 0.7). This suggests that a substantial component of the masking associated with 4-talker babble derives from acoustic-linguistic similarities between the target, which is often referred to as informational masking (56, 60, 71). The consistency of these results with prior literature confirms the viability of the online testing platform in reproducing in-person measurements.

Figure 5 (b) illustrates masking release for the same four conditions as in Figure 5 (a), except for the addition of reverberation in all conditions. Note that the reference condition (i.e., babble speech with no F0 or spatial cues) also contained reverberation. Reverberation generally reduced the masking-release benefit, except for the F0-only condition. This is consistent with previous studies showing that reverberation has a smaller impact on the use of monaural cues (54, 72, 73), while spatial hearing is subject to substantial degradation (36, 54, 74).

**Fig. 5.**
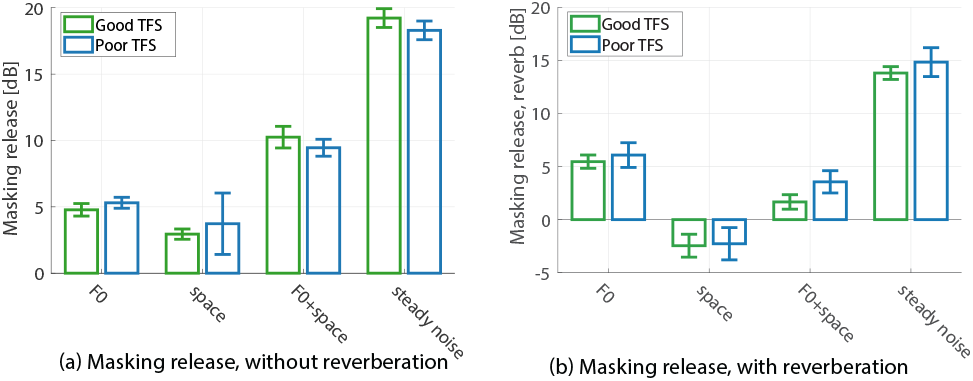
Masking release across conditions. The height of the bars represents the mean, error bars represent +/-1 std. Masking release was calculated by subtracting the SRT in each condition from that for the reference condition. Note that the reference condition in (a) does not have reverberation, whereas the reference condition in (b) contains reverberation. A positive masking release means that the SRT was lower/better than that for the reference condition.

Figure 5 (a) and 5 (b) demonstrate similar masking release for participants divided into two groups based on their TFS sensitivity. In both non-reverberant [Figure 5 (a)] and reverberant conditions [Figure 5 (b)], the Good-TFS group did not benefit more from the cues in terms of masking release in any of the conditions tested. This is consistent with other studies suggesting that better TFS processing might not necessarily benefit a listener by conferring *more* masking release when envelope-based cues are also available (17, 35, 63, 75).

### Better TFS processing is associated with resilience to the effects of reverberation and reduced listening effort for speech perception in noise

To illustrate the advantage associated with robust TFS processing for listening under reverberation, the threshold increase from non-reverberant to reverberant conditions is shown by the height of the bars in Figure 6 (a). The group with poor TFS sensitivity (*mean* = 5.5, *std* = 0.4) showed a greater threshold increase in reverberant settings than their good-TFS counterparts (*mean* = 3.2, *std* = 0.3) (Figure 6 (a), left; *z* = 4.6, *p* = 0.2*e−*4). When the participants were divided based on their ILD sensitivity, there was no significant group difference, indicating an important role for TFS processing. This result suggests that better TFS sensitivity can mitigate the negative impact of reverberation, which is a common source of signal degradation in everyday listening environments.

**Fig. 6.**
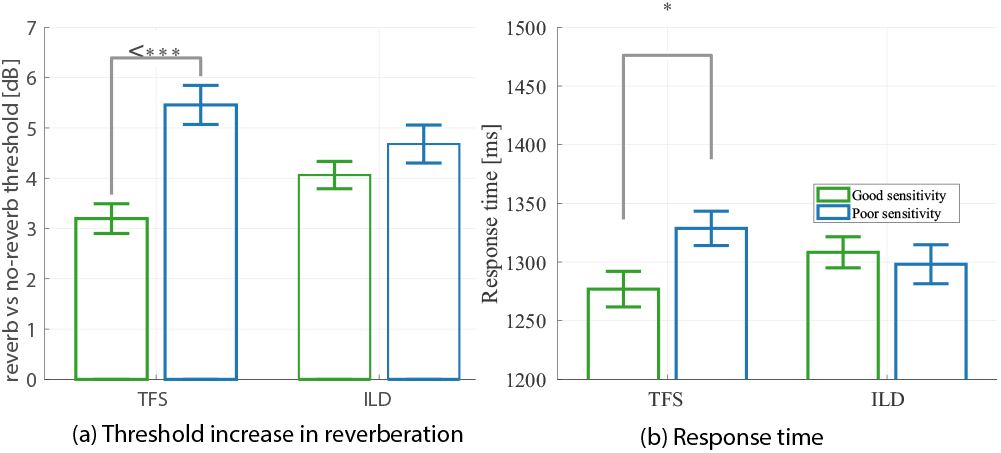
(a) Increase in SRT due to reverberation for each group of participants. (b) Average response times. Data were pooled across all conditions shown in Figure 5. All error bars represent estimated standard error of the mean. Significance stars: .05 *> * ≥* .01, .001 *> * * * ≥* .0001 (corrected for multiple comparisons using FDR procedures).

It is well known that behavioral measures of performance may not reveal important differences in the cognitive effort expended by participants in achieving a given level of performance (76, 77). To investigate whether robust TFS sensitivity is associated with less effortful listening, we examined response times, a measure commonly utilized in the literature for assessing listening effort (61, 62, 78, 79). The response times are indicated by the height of the bars in Figure 6 (b). The absolute values of the response times are consistent with prior literature (80). When the participants were divided into two groups based on their TFS sensitivity, the Good-TFS group (*mean* = 1277, *std* = 15.2) exhibited significantly shorter reaction times than the Poor-TFS group (*mean* = 1328, *std* = 14.6) (Figure 6 (b), left; *z* = *−*2.5, *p* = 0.035), consistent with reduced listening effort for the former. When the participants were regrouped based on non-TFS characteristics (i.e., ILD sensitivity), there was no significant difference between the two groups (Figure 6 (b), right). Taken together, these results show that robust TFS sensitivity is associated with shorter reaction times. Both of these results, i.e., the smaller decrement in performance under reverberation and smaller overall response times in the good TFS group, remain significant after correcting for multiple comparison (10 comparisons across Figures 5 (a) and (b), 6 (a) and (b) using false discovery rate (FDR) procedures (81) at a 5% FDR level.

### New measurements replicating the study corroborate the main findings from the original experiments

As suggested by an anonymous reviewer, we reached out to all 200 individuals who participated in 2020. Given the intermission of 3+ years, there was substantial attrition. Forty four participants responded and completed the replication measurements. The replication experiments were more narrowly focused to test the main claims from the original study. Specifically, the measurements included the measures and controls used for grouping (i.e., ITD, binaural FM, and ILD), and speech-in-noise measurements in anechoic and reverberant settings. Because the goal was to test the effects of reverberation, we only included the reference and F0-cue conditions. Despite a gap of more than three years, we observed statistically significant correlations between the original and repeated TFS-sensitivity measurements (ITD and binaural FM measurements, Figure 7, A2 and A3). With the same grouping method being applied to the replication dataset for TFS-sensitivity measures (Figure 7, B1), we see similar results as in Figure 6: smaller increase in speech reception thresholds due to reverberation (Figure 7, B2) and shorter response time (Figure 7, B3) overall for the Good-TFS group. When only the top and bottom 25% of the replication sample were chosen for grouping (Figure 7, C1), to increase the group difference in TFS sensitivity, the corresponding differences in the reverberation effects and response times also increased (Figure 7, C2 and C3). Although not the focus of the replication study, note that F0-based masking release was not significantly different between groups (for groups in Figure 7 B1 and C1), which is consistent with the original results from Figure 5. In summary, despite the delay between the original and the replication experiments, the new data corroborated both key findings from the original study and provided further credence to the notion that binaural measures can robustly capture individual differences in TFS processing.

**Fig. 7.**
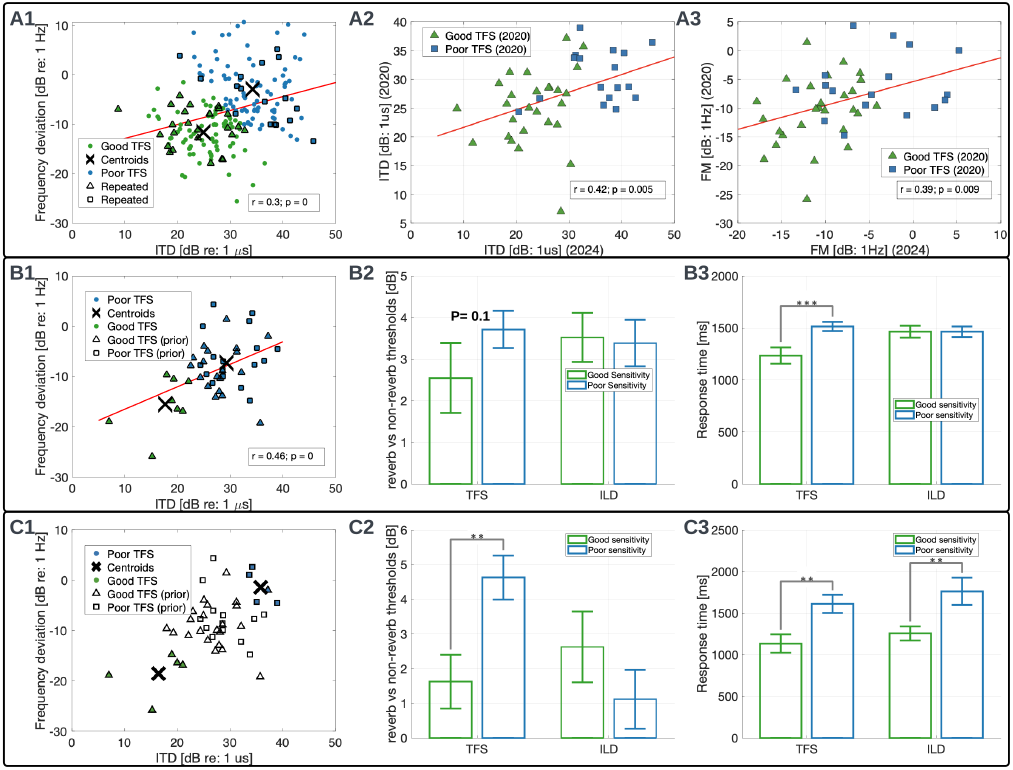
Summary of the replication results obtained in 2024 from a subset of the same cohort. A1-A3: there are statistically significant correlations between the original and replication data; B1-B3: the Good TFS-sensitivity group, based on replication measures, showed less increase in SRT due to reverberation and shorter response time overall. C1-C3: group differences in the reverberation effects and response times increased with greater group difference in TFS sensitivity. Significance stars:.01 *> ** ≥* .001, .001 *> * * * ≥* .0001 (corrected for multiple comparisons using FDR procedures).

## Discussion

No *greater* spatial release from masking was observed for the “Good-TFS” group despite the theoretical connection between TFS phase locking and binaural temporal processing (21, 22) [Figure 5 (a)]. Brainstem binaural circuits compare temporal information encoded by TFS phase locking from each ear and can encode microsecond ITDs that form one of two main cues supporting spatial hearing along the horizontal plane. Accordingly, we hypothesized that individuals with better TFS sensitivity would benefit more from the spatial cues in speech-in-noise tasks. One of the reasons why we did not find a group difference may be that the participants were all typically hearing; individual differences in TFS sensitivity may not have been sufficiently large. A group difference may be observable if a broader range of TFS sensitivity is represented in the cohort by including individuals with hearing loss. Similar to our finding, Füllgrabe et al., 2015 (63) did not observe an age effect on spatial release from masking, which might have been limited by a smaller age effect on TFS sensitivity from their typical hearing older participants. Another plausible reason could be that the spatial cue in this study was large (i.e., *S*_π_ *N*_0_ vs. *S*_0_*N*_0_). There might have been a group difference for a small ITD difference between target and masker. Finally, the use of ILD discrimination as a reference for non-TFS factors could also have contributed to the lack of group difference in spatial release from masking. ILDs also activate binaural circuits, although ILD-based binaural processing does not rely on TFS phase locking (82). Regressing out ILD scores from binaural TFS measurements could have removed any individual variability in aspects of spatial hearing that go beyond sensitivity to TFS cues, such as the efficacy with which downstream “readout” processes use binaural information. Thus, rather than contradicting the prevailing view that TFS processing is critical to spatial hearing (7, 21, 22, 38, 83, 84), our result simply suggests that the range of individual differences observed in ITD thresholds did not translate to measurable differences in the degree of spatial release from masking.

Similarly, no significant group difference was observed for F0-based masking release. Although TFS processing is widely acknowledged as important for low-frequency spatial hearing, its role in pitch perception has been debated for over 150 years (85, 86). Humans perceive low-frequency periodic sounds as having a stronger pitch than high-frequency sounds (23–25). Frequency discrimination threshold, expressed as Δ*F/F*, increases with increasing frequency from 2 to 8 kHz, plateauing above 8 kHz (87–89), which aligns with the low-pass characteristic of TFS phase locking in the auditory nerve (90, 91). Deficits in TFS coding have been invoked to explain speech perception deficits in fluctuating noise (41), where target-masker F0 differences are thought to play a role (92, 93). While these findings appear to suggest that TFS may play a role in pitch perception, the same observations also permit alternative interpretations based on place coding, which also worsens at higher frequencies and in individuals with hearing loss (14, 94). The result from the present study, i.e., the similar F0-based masking release across Good- and Poor-TFS groups leans towards “place-coding” based explanations of pitch phenomena.

The “steady noise” condition used in the present study [Figure 5 (a)] was designed to minimize modulation masking (interference from modulations in the maskers) so that energetic masking would be dominant (95) (see Methods). In contrast, the 4-talker babble masker in the reference condition contained many sources of modulations and informational masking (e.g., modulation masking, phonetic/lexical/semantic content) in addition to energetic masking (96). The improvement of almost 20 dB in SRTs from the reference to the “steady noise” condition [consistent with Arbogast et al., 2002 (71)] points to the dominant role of informational masking in everyday listening (97). Listening in the presence of informational masking is thought to involve many sensory and cognitive processes in the central auditory system, including object formation and scene segregation/streaming, auditory selective attention, working memory, and linguistic processing (55, 56). TFS-based processing is thought to play an important role for scene segregation and attentive selection (58–60). Although our results show similar release from informational masking across the two TFS-sensitivity groups [Figure 5 (a)], the group with better TFS sensitivity had a significantly shorter response time than the poorer TFS group [Figure 6 (b)]. Our results, therefore, affirm the contribution of TFS coding to robust central auditory processing, possibly with lower listening effort. The fact that the group difference in reaction times did not translate into the masking-release metrics underscores the need to investigate cognitive factors beyond performance/score metrics to fully characterize the importance of different peripheral cues (98–100).

Finally, we explored the correlation between TFS processing and listening in a reverberant environment. The SRTs were considerably worsened by the presence of reverberation [Figure 6 (a)]. More importantly, the group with poor TFS sensitivity was affected significantly more than their good-TFS counterparts, indicating a possible role of TFS processing in resisting the deleterious effects of reverberation. Reverberation impairs TFS-based spatial cues (36) and spatial selective attention (54). Thus, our findings suggest that stronger TFS coding may ameliorate reverberation’s detrimental effects on speech perception in noise.

These observations, together with the fact that most cochlear implants (CIs) do not convey TFS also help explain the effortful listening experience of CI users, especially in the presence of reverberation. The findings also suggest that evaluation of TFS processing may complement conventional assessments used in audiology clinics to help characterize speech perception deficits in background noise (54, 101, 102). Although the combined use of ITD, binaural FM and ILD measures shows potential for capturing individual differences in TFS sensitivity, further validation and refinement is needed before they can be feasibly applied to clinical settings. Finally, our results also affirm the promise of using web-based psychoacoustics to conduct large-scale experiments (50, 53). Automated data collection facilitates the rapid acquisition of data from a large participant cohort over a short time frame (several days), providing a substantial advantage over traditional in-person psychoacoustic testing. Finally, whether the perceptual benefits associated with better TFS sensitivity directly derive from the TFS code, or whether both derive from other common physiological factors, cannot be ascertained in this study. Although the contribution of non-sensory variables such as motivation and attention was mitigated by using the ILD metric as a control (51), there may be factors that preferentially affect the TFS code while also affecting speech in noise through mechanisms distinct from TFS processing. One such candidate mechanism is cochlear neural degeneration, which is hypothesized to affect temporal coding (48), and can also trigger central auditory changes which in turn can impair listening in the presence of background noise (103, 104).

## Materials and Methods

### Participants

Two hundred participants were recruited anonymously from Prolific.co (20-55 years old (mean=31, std=8); 93 females, 102 males, and 5 not reported). Eighty five percent of the participants self-reported English as their first language, and all participants were native speakers of North American English. In terms of race and ethnicity, 64% self reported as White, 21% as Asian, 6.5% as Mixed, 2.5% as Black, 2.5% as Other, and 3.5% not reported. Participants reported no hearing loss, neurological disorders, or persistent tinnitus, and passed headphone checks and a speech-in-noise-based hearing screening (53). The participants consented to participate following Institutional Review Board (IRB) protocols established at Purdue University and were compensated for their time. All participants completed the full study battery. The median time for completion was approximately 1 hour.

### Experimental Design and Statistical Analyses

#### Screening Measurements

All measurements, including the screening, are listed in Figure 8. Because participants were anonymous and used their own computers and headphones, two screening procedures were administered to narrow the pool of participants to individuals with typical hearing, and to ensure stereo headphone use.

**Fig. 8.**
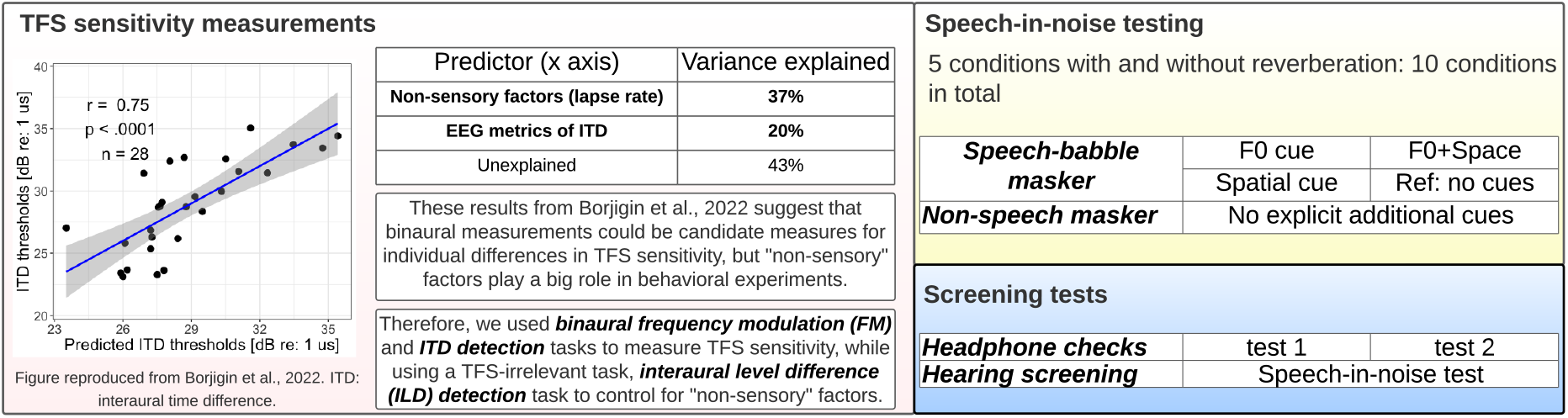
An illustration of all measures included in the present study.

#### Headphone-Check

Two tests based on previously established procedures were carried out to screen for appropriate use of headphones (53). In the first, participants were instructed to identify the softest of a sequence of three low-frequency tones. The target tone was 6 dB softer than the two foil tones, but one of the decoy tones was presented with opposite phase at the left and right channels (105). Woods et al., 2017 (105) reasoned that if a participant used a pair of sound field loudspeakers instead of headphones, acoustic cancellation would result in an attenuation of the anti-phase decoy tone leading to an error. However, the procedure becomes ineffective if a participant uses only one loudspeaker/channel. To catch participants who used a single-channel set up, we added a second task where participants were asked to report whether a low-frequency chirp (150–400 Hz) embedded in background low-frequency noise was rising, falling, or flat in F0. The stimulus was designed such that chirp was at pi-phase between the left and right channels, whereas the noise was at zero phase (i.e., a so-called “S*π*N0” configuration). The signal-to-noise ratio was chosen such that the chirp would be difficult to detect with just one channel, but easily detected with binaural headphones because of the so-called binaural masking level difference (106).

#### Hearing Screening

Participants were screened for hearing status using a speech-in-noise task previously validated for this purpose (53). A previous meta-analysis of 15 studies suggested that speech-in-noise tasks yield a large effect size, separating individuals with typical hearing and hearing loss, and can thus serve as sensitive suprathreshold tests for typical-hearing status (53). A speech-in-babble task was administered to a cohort of individuals with known hearing status (either audiometrically typical hearing or known degree of hearing loss) and cutoff values were chosen based on the scores obtained such that the procedure yielded *>* 80% sensitivity to any hearing loss, and *>* 95% sensitivity to more-than-mild hearing loss (53). Together with the headphone-check procedure, the speech-in-noise hearing screening helped narrow the pool of participants to those who used 2-channel headphones, had typical hearing, and were in good compliance with the study instructions. Two hundred participants who passed all screening procedures proceeded with the main battery of the study. No training was provided except for a brief demonstration block for each task.

#### TFS Sensitivity Measurements

We previously established that binaural behavioral and electrophysiological (EEG) measurements of ITD sensitivity can reliably reflect individual differences in TFS processing (51). Therefore, in this study, we adopted behavioral ITD detection and added a binaural version of frequency-modulation (FM) detection. Importantly, our previous study also showed that the binaural metrics were effective in capturing individual differences in TFS processing only if the contributions of extraneous “non-sensory” factors that are irrelevant to TFS processing, such as engagement, were measured and adjusted for (51). In the present study, we implemented a stand-alone measure that would also be influenced by extraneous non-sensory factors, but unaffected by TFS processing. Specifically, we used an interaural level difference (ILD) discrimination task, which is also a binaural task but depends on level coding instead of TFS coding. The use of ILD discrimination as a surrogate measure not only helped mitigate non-sensory extraneous variability, but also likely enhanced the specificity of the ITD and binaural FM measures to TFS processing by removing individual variability in downstream “readout” processes that used binaural information.

#### Interaural Time Difference (ITD) Discrimination

The stimulus consisted of two consecutive 400-ms-long, 500-Hz pure tones. The tones were delivered to both ears, but with a time delay in one randomly selected ear (i.e., ITD). The leading ear was switched from the first to the second tone in the sequence, simulating a spatial “jump” to the opposite side. ITDs in steps of a factor of two from 2 to 128 *µs* were presented in random order (8 repetitions for each step). The tone bursts were ramped on and off with a rise and fall time of 20 ms to attenuate abrupt stimulus-silence transition and to reduce reliance on onset ITDs. The gap between the two tone bursts was 200 ms. As with other tasks, participants were instructed to adjust the volume control on their devices to a comfortable loudness. A two-alternative forced-choice (2AFC) task was used, where participants were asked to report the direction of the “jump” between the two intervals (left-to-right or right-to-left) using a mouse click. A separate “demo” block was provided before the experimental blocks to familiarize the participant with the task. The detection thresholds were quantified using a Bayesian approach (107, 108), using the psignifit toolkit from wichmann-lab. The same method for estimating thresholds was used for all measurements of this study, including TFS and ILD sensitivity, and speech-in-noise measurements (Figure 8).

#### Binaural Frequency Modulation (FM) Detection

We employed a binaural FM detection task as a second metric of individual TFS sensitivity. Although low-rate monaural FM detection has been used to probe TFS processing (102, 109, 110), whether monaural FM detection can truly measure individual TFS processing fidelity is questionable (49, 51). In contrast, binaural temporal processing has an unambiguous theoretical connection to TFS coding (22, 51). The binaural FM detection measure implemented in the present study consisted of target and reference stimuli in a 2AFC task. The stimuli in each interval were turned on and off with a rise and fall time of 5 ms to attenuate abrupt stimulus-silence transition. The reference was a 500-ms, 500-Hz diotic pure tone. The target tone had a 2-Hz rate FM around 500 Hz with modulation out of phase in two ears to introduce binaural timing cues. A low FM rate was chosen because of the “sluggishness” of binaural system: our inability to track fast binaural modulations (111, 112). FM depths (maximum frequency deviation in one direction) in steps of a factor of 2 from 0.1 to 3.2 Hz were presented in random order (8 repetitions for each step). The starting phase of the stimuli was set at 0. No training was provided except for a brief demonstration block that was intended for orienting the participants before the formal testing.

#### Interaural Level Difference (ILD) Discrimination

ILD discrimination thresholds were measured with two consecutive 4-kHz pure-tone bursts, a frequency where TFS phase locking is generally thought to be limited (5). Similar to the ITD task, the two intervals were lateralized to opposite sides through ILDs, simulating a spatial “jump” from one side to the other. ILDs in steps of a factor of 2 from 0.1 to 3.2 dB (8 repetitions for each step) were presented in random order. Participants were asked to report the direction of the jump through a mouse-click response in a 2AFC task. A similar approach was used by Flanagan et al., 2021 (113), where they used intensity discrimination as a covariate in the statistical analysis to control for monaural factors since the study’s focus was binaural processing. In this study, since we used binaural measurements as TFS sensitivity measures although binaural processing itself is not the focus, we used interaural level difference discrimination to also control for the binaural factors.

#### Rationale

The TFS (ITD and Binaural FM) and control (ILD) measures, and sample size (n = 200) chosen here were guided by findings from our previous study showing robust EEG-behavior correlations in TFS measures with about 40 participants (51). However, that was an in-person study. Because the variance across participants in web-based measures is generally about 75-90% larger with our platform [see Table 1 in Mok et al., 2023 (53)], we doubled the participant number and did so for each group (effectively quadrupling the sample size for individual difference comparisons).

#### Grouping of Participants

Participants were classified into two groups (Good vs. Poor sensitivity), either based on TFS-sensitivity measures or the ILD measure [see Figure 2 (a) and (b)]. A two-dimensional “k-means” clustering algorithm was used for grouping based on the two TFS measures whereas a simple median split was used for ILD-based grouping (given that it was based on a single measure). Note that, before clustering, ILD sensitivity was “regressed-out” from the two TFS-sensitivity measures using a simple linear regression to emphasize individual differences in TFS processing and mitigate the effects of extraneous variables on the TFS measures. Although ILD detection is supposed to be more or less independent of TFS processing, it is subject to non-sensory contributions from variables like attention/motivation etc. that can introduce spurious correlations between ILD detection and speech-in-noise. The mutual “regressing out” of ILD and TFS measures from each other can help reduce these non-sensory contributions.

#### Measurements of Speech Perception in Noise

The stimuli consisted of a target word with a carrier phrase (Modified Rhyme Test) and a masker. The masker was either four-talker babble (IEEE speech corpus) or a steady noise composed of an inharmonic complex of tones (95), described below. The carrier phrase was in the same voice as the target word and said: “Please select the word …”. The masker began after the onset of the target carrier phrase but before the target word to allow participants to orient themselves to the target voice based on the unmasked portion of the carrier phrase. A word-based test rather than a sentence-based test was chosen to minimize the influence of factors such as individual differences in working memory, and ability to use linguistic context.

Participants were tested across 10 target-masker conditions, as shown in Figure 8. Four conditions used four-talker babble as the masker and one used a non-speech, steady masker. The babble masker conditions included F0 cues, spatial cues, both F0 and spatial cues, and no explicit cues (i.e., reference). Note that the 4-talker babble consists of speakers of the same sex. For conditions with F0 cues, if the target was a male talker, for example, the 4-talker babble would consist of female talkers. The non-speech masker condition had a steady masker without any explicitly added cues. The remaining 5 conditions were similar but with the addition of room reverberation. The presentation order of the 10 test conditions was randomized across trials. Details about the stimulus manipulations used are provided below. For each condition, speech intelligibility was measured over a range of SNRs to estimate the speech reception threshold (SRT), defined as the SNR at which approximately 50% of the words were intelligible.

#### F0 Cues

To control the available F0 cues for separating the target and masker, the audio recordings for all trials were first processed to remove inherent F0 fluctuations (i.e., monotonized to the estimated F0 median) using Praat (version 6.4.04) and a custom Praat script (written by Matthew B Winn). Then, the flattened F0 contours of each target sentence and each talker in the four-talker babble were transposed to a preset value, as shown in Figure 9. The F0 of female target voice was set to 245 Hz, and that of the male target was set to 95 Hz. Among the talkers whose sentences were mixed to create the four-talker babble background, the male talkers’ F0 values were set to 85, 90, 100, and 105 Hz, and the female talkers’ F0 values to 235, 240, 250, and 255 Hz. Note that the target and masker of the same sex had similar F0 values but with a small difference to ensure that the participant could still distinguish the target from the masker but could only derive minimal masking release based on F0 difference. The F0 contour was flattened for all other stimulus configurations (i.e., reference, space, F0+space, and non-speech noise masker). F0-based masking release was estimated as the SRT difference between the reference condition where the target and masker stimuli were composed of recordings from same-sex talkers and the “F0” condition where there was a large F0 separation by virtue of the target and masker stimuli being composed of recordings from opposite-sex talkers.

**Fig. 9.**
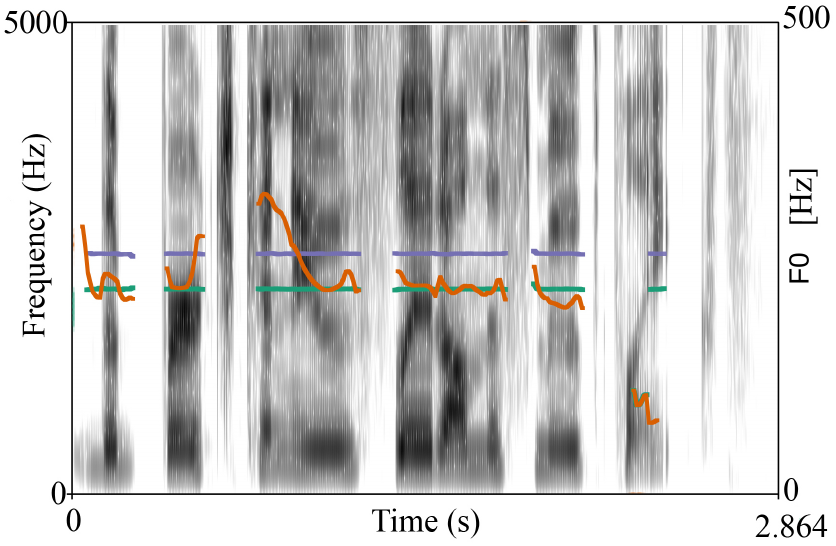
The spectrogram of a sentence: “The birch canoe slid on the smooth planks.” The orange curve shows the estimated F0 contour with natural fluctuations; the flattened F0 contour is shown in green; the flattened F0 contour that was transposed to a pre-set frequency (255 Hz in this example), is shown in purple.

#### Spatial Cues

To simulate the perception of spatial separation using purely TFS-based cues, the polarity of the target in one ear was inverted while the masker was kept the same in the two ears. This configuration is denoted *S*_*π*_*N*_0_. The fully diotic condition without this interaural manipulation is referred to as *S*_0_*N*_0_. A lower SRT (i.e., better performance) is typically observed in the *S*_*π*_*N*_0_ condition, the difference in SRTs denoted the binaural masking level difference (BMLD, i.e., spatial release from masking) (114).

#### Steady Masker and Reverberation

Performance in the presence of a steady masker was used to evaluate the role of TFS in providing release from so-called “informational masking” (96). Accordingly, the steady masker was designed to have minimal intrinsic modulations and match frequency content to the typical speech frequency range (1-8000 Hz) using the procedure described in (95). The masker was dichotic, consisting of odd-numbered sinusoids delivered to one ear and even-numbered sinusoids to the opposite ear. This approach reduced the occurrence of beats generated by neighboring components in the peripheral auditory system, ensuring minimal amplitude fluctuations of the masker at the outputs of the auditory filters. Owing to the lack of modulations (explicit and intrinsic), this masker represents a condition where energetic masking is dominant while avoiding most sources of informational masking. Note that conventionally used noise maskers such as speech-spectrum stationary noise have intrinsic modulations that can contribute to masking at more central levels of the auditory system (97, 115–117). Finally, to simulate listening under reverberation, the stimuli that were recorded under anechoic conditions were convolved with binaural room impulse responses recorded in a bar (BarMonsieurRicard.wav from echoThief).

#### Speech Reception Threshold (SRT) Estimation

To robustly estimate the mean and variance of the masking release based on different cues, SRTs for each speech-in-noise condition were estimated using a jackknife resampling procedure. Within each group (Good vs. Poor), a leave-one-out procedure was used: psychometric functions were fit to the percent-correct vs. SNR scores that were obtained by averaging the data across all participants except the one being left out. The SRT was then estimated as the midpoint of this psychometric curve. Across individuals within a group, this procedure generated *k* jackknife samples for the SRT for each condition and masking release for each cue (where *k* is the number of individuals within the group). Following (118), the group-level mean *M* was estimated as the mean across the jackknife samples, and the variance as the sample-variance *V* across the jackknife samples multiplied by (*k −* 1). The jackknife procedure avoids the need to fit psychometric curves for speech intelligibility as a function of SNR or to estimate SRTs at the level of the individual participant, and yet robustly estimates the variance in the SRTs (and masking release values) across participants within each group. ***Response Time***. Two participants with comparable SRTs could experience different levels of listening effort (76, 77). To assess the role of listening effort, the reaction time for each participant was determined by subtracting the time of the stimulus offset (or stimulus duration) from the recorded time of the mouse-click response. The same procedure as for the SRT estimates was used to estimate the mean and variance of the response times. Trials with response times larger than 10 seconds were discarded, under the assumption that they were likely due to interruptions in participation rather than the engagement of cognitive processes to select a response. Response times were separately estimated for each participant group, and for each speech-in-noise condition.

#### Statistical Analyses

The primary analyses involved between-group comparisons of masking release or response times. Because the cohort size was large (N=200) and estimates of group mean and variance were derived using the jackknife procedure, it was reasonable to assume that group-level estimates represented parameter estimates for normally distributed data. Accordingly, simple one-tailed z-tests were used for making inferences. As described previously, among the 10 speech-in-noise conditions, 5 simulated speech-in-noise mixtures in anechoic environments and 5 included room reverberation. To investigate the effects of reverberation, data from all 5 speech-in-noise configurations were combined using inverse variance pooling (119, 120). For response time comparisons across groups, all 10 conditions were pooled.

## Data Archiving

The data, scripts for setting up online experiments, data analyses, and step-by-step instructions have been uploaded on the Open Science Framework (OSF project) and will be made available upon publication.

## ACKNOWLEDGMENTS

We thank our research participants for their participation in this study. This research was supported by funding from the National Institutes of Health [Grant R01DC015989]. Portions of the paper were developed from the thesis of A.B. (121).

